# A major 6 Mb superlocus is involved in pyrethroid resistance in the common bed bug *Cimex lectularius*

**DOI:** 10.1101/2023.01.06.522975

**Authors:** Chloé Haberkorn, Jean-Philippe David, Hélène Henri, Jean-Marie Delpuech, Romain Lasseur, Fabrice Vavre, Julien Varaldi

## Abstract

In the last few years, the bed bug *Cimex lectularius* has been an increasing problem world-wide, mainly due to the development of insecticide resistance to pyrethroids. The characterization of resistance alleles is a prerequisite to improve surveillance and resistance management. To identify genomic variants associated with pyrethroid resistance in *Cimex lectularius*, we compared the genetic composition of two recent and resistant populations with that of two ancient-susceptible strains using a genome-wide pool-seq design. We identified a large 6 Mb “superlocus” showing particularly high genetic differentiation and association with the resistance phenotype. This superlocus contained several clustered resistance genes, and was also characterized by a high density of structural variants (inversions, duplications). The possibility that this superlocus constitutes a resistance “supergene” that evolved after the clustering of alleles adapted to insecticide and after reduction in recombination is discussed.

## 1 INTRODUCTION

The common bed bug *Cimex lectularius* (Hemiptera) is a blood-sucking human parasite that can cause both physical and psychological disorders, such as allergic reactions (Alexander, 1984), sleep deprivation (Susser et al., 2012) or even paranoia (Goddard and Deshazo, 2009). Although the massive use of insecticides in the 20th century helped to regulate their populations, a phenomenon of resurgence of the species has been observed since the late 1990s (Potter, 2011). This population boom in bed bugs is thought to be due to the increase in international travel, the growing popularity of second-hand market (Doggett et al., 2004), and the spread of insecticide resistance (Davies et al., 2012). Pyrethroid insecticides are widely used against bed bugs, thereby exercising a strong selection pressure. Hence, the current rise of bed bugs is believed to be mainly imputable to the emergence of pyrethroid resistance (Romero et al., 2007).

Insecticide resistance mechanisms have long been studied in a variety of insects, and are divided in two types: behavioral resistance, and physiological resistance (Lockwood et al., 1984). Behavioral resistance is defined as the ability to avoid or reduce lethal exposure to an insecticide. Physiological resistance consists in decreasing insecticide intake by preventing its entry, with decreased cuticular penetration (Balabanidou et al., 2018), or limiting its effect, through increased detoxification metabolism and decreased target site sensitivity (Feyereisen, 1995).

To date, no convincing evidence of behavioral resistance has been found in *C. lectularius* (see review Dang et al. 2017), although some behaviors that may limit the effect of the insecticide have been observed in both resistant and susceptible strains. Pyrethroid insecticide could indeed have an excitatory-repulsive effect on bed bugs, with increased movement whenever in contact, or the choice of an untreated place to hide (Romero et al., 2009a).

Cuticular resistance can be the consequence of a thickening or change in cuticular structure that decreases the rate of insecticide penetration. A morphological study confirmed that cuticle was thicker in a pyrethroid resistant strain of bed bugs collected in Australia, compared to a susceptible one (Lilly et al., 2016b). An increased cuticle thickening was correlated with a longer time-to-knockdown after insecticide exposure. Further studies based on RTqPCR assays showed that cuticular protein genes, such as chitin synthase (CHS) or larval/pupal cuticle protein (L/PCP), had higher transcript levels in resistant bed bug strains (Mamidala et al., 2012; Koganemaru et al., 2013).

Following penetration of the cuticular barrier, detoxification metabolism may be mobilized to degrade, sequester and enhance the excretion of the insecticide. In resistant populations of bed bugs, an increased activity of several detoxification enzymes such as cytochromes P450s (Romero et al., 2009b) and esterases (Lilly et al., 2016a) has been observed. Higher expressions of glutathione-S-transferases (GSTs) have also been detected in resistant bed bug immature stages, but this mechanism is yet to be identified in adults (Mamidala et al., 2011). Some studies also used specific enzymatic inhibitors to determine the involvement of particular detoxification enzyme families in the resistant phenotype. Piperonyl butoxide (PBO), an inhibitor of P450s, significantly decreased resistance against the pyrethrinoid deltamethrin (Romero et al., 2009b). Similarly, an inhibitor of esterases increased bed bug mortality to the pyrethroid lambda-cyhalothrin (Lilly et al., 2016a).

Pyrethroid being the main chemical insecticide used today against bed bugs, its target protein site is also particularly studied (Dang et al., 2014, 2015; Akhoundi et al., 2015; Balvín and Booth, 2018). Pyrethroids target the voltage-gated sodium channels (*VGSC*) in the nervous system of the insect. When pyrethroids bind to the *VGSC*, they induce paralysis of the insects by maintaining the channels in an open state (knock-down effect). A single nucleotide non-synonymous mutation can alter *VGSC* conformation, and therefore hinder pyrethroid binding, thus conferring knock-down resistance (*kdr* mutations).

Three *kdr* mutations have been associated with pyrethroid resistance in *C. lectularius*: V419L (valine in position 419 to leucine), L925I (leucine in position 925 to isoleucine), and I936F (isoleucine at position 936 to phenylalanine). V419L and L925I *kdr* mutations were detected in Eurasia and USA in early 2000s, while the I936F mutation was detected in Australia during the same period (Zhu et al., 2010, 2013; Dang et al., 2015). According to Durand et al. (2012), all bed bugs sampled in Paris (n=124) during 2011 were homozygous for the L925I mutation.

In bed bugs, many reports showed resistance through phenotypic assays to various insecticides (Romero and Anderson, 2016; Lilly et al., 2018; Cáceres et al., 2019). Using knowledge obtained on other insect systems, the association between resistance phenotype and genes has been explored for a few candidate genes, such as *VGSC* (Zhu et al., 2010). Other wider approaches identified genes potentially involved in resistance by comparing the transcriptomic profile of resistant and susceptible lines (Adelman et al., 2011; Mamidala et al., 2012).

In an attempt to further identify the underlying genetic factors of resistance, Fountain et al. (2016) went through a RAD-tag based QTL mapping of resistance to deltamethrin by crossing a susceptible strain with a pyrethroid resistant one, both originating from London. Using 334 SNP markers, they found a major QTL associated with insecticide resistance on a Linkage Group (LG 12). Interestingly, this 10 Mb region contains several genes considered as valuable candidates for pyrethroid resistance, such as the *VGSC*, a carboxylesterase and several P450s. However, identified genes alone failed to fully explain variations in resistance phenotype, with the major QTL detected explaining only 64.2% of phenotypic variation. Moreover, since this study was based on RAD-tags only, it did not provide the full sequences of these putative resistance variants. Nor did this analysis provide any information on structural variants. Indeed, structural variants such as duplications and inversions have been shown to be potentially involved in insecticide resistance, as in the mosquito *Anopheles gambiae* (Brooke et al., 2002; Assogba et al., 2015). Inversions can inhibit recombination between standard and inverted regions in the heterozygote state, thus having a potential protective effect on a large number of genes (Ayala et al., 2014; Mérot et al., 2020). Other structural variants, as gene copy number, could also lead to insecticide resistance. Changes in gene copy number (Copy Number Variation, or CNV), and specifically duplications, can lead to overexpression of genes through a “gene-dosage” effect (Kondrashov, 2012). Amplification of genes encoding detoxification enzymes can enhance insecticide metabolism in many insects, and most notably via three gene families: esterases, glutathione S-transferases (GSTs), and cytochrome P450 monooxygenases (P450s) (Bass and Field, 2011). However, the role of structural variants in the capacity of bed bugs to resist insecticides has not yet been investigated.

In this study, we aim to provide a genome-wide analysis of the genetic factors underlying deltamethrin resistance in bed bugs. We performed DNA pool-sequencing on four strains of *C. lectularius* with different insecticide resistance statutes. Based on both single nucleotide polymorphisms (SNPs) and structural variants (SVs) detection, we used population genomics to identify allelic variants and genomic regions associated with the resistance phenotype.

## 2 MATERIALS AND METHODS

### 2.1 Insects

The four strains used in this study were provided by CimexStore Ltd (Chepstow, United Kingdom). Two of these strains were susceptible to pyrethroids (S), as they were collected before their massive use and have been maintained under laboratory condition without insecticide exposure for more than 40 years: German Lab (GL, collected in Monheim, Germany) and London Lab (LL, collected in London, Great Britain). The other two resistant (R) populations were London Field (LF, collected in 2008 in London) moderately resistant to pyrethroids, and Sweden Field (SF, collected in 2015 in Malmö, Sweden), with a moderate-to-high resistance level, according to CimexStore statement. These statutes were confirmed by our bioassays (see below).

Insects were kept isolated before imaginal moult in 24-wells Petri dishes containing accordion-folded blotting papers, serving as harborage. Bed bugs were maintained at 25°C, 40% relative humidity (RH), and a photoperiod of 12:12h. Since males perform the so-called traumatic insemination to copulate, which can be costly for their partners (Stutt and Siva-Jothy, 2001), bioassays were conducted only on 7-day-old unfed virgin females. Insecticide resistance is thought to be autosomal in bed bugs (Feroz, 1969; Fountain et al., 2016).

### 2.2 Topical Assay

The insecticide resistance level of the “field” strains and the susceptibility of the “lab” strains was first confirmed by determining the resistance ratio between these strains. Topical insecticide bioassays were carried out with deltamethrin (98% purity, Cluzeau, Sainte-Foy-La-Grande, France), a pyrethroid, as it remains the most used insecticide family recently for bed bug control. Three replicates of 10 insects were used both for control and per insecticide dose, ranging from 0.01 ng to 120 ng per µL. Bed bugs were previously immobilized by placing them in a Petri dish on ice for 5 minutes. Topical applications were then made onto the ventral surface of the thorax, between the coxae, with a 50-µL glass syringe attached to a repeating dispenser (Hamilton Co., Reno, NV). Treated insects were exposed to 1 µL of insecticide diluted in acetone, whereas control insects received 1 µL of acetone only. Mortality was assessed after 24 h by flipping each insect on dorsal side with a featherweight forceps, to see if it was able to reverse on the ventral side (alive) or if its movements were not coordinated enough to do so (moribund, considered as dead).

Dose–mortality data were analyzed by using binomial Generalized Linear Models with a probit transformation using the R/ecotox package v1.4.4 (Hlina et al., 2021). Statistical tests and resistance ratios were computed on lethal doses for 95% of individuals (LD_95_). Resistant ratios were calculated on pairs as follows: highest LD_95_ (i.e. of the more resistant population of the pair) divided by the lowest LD_95_ (more susceptible population).

### 2.3 DNA extraction and sequencing

For each strain, genomic DNA was extracted from 30 individual females (except for London Lab which had only 28) using NucleoSpin 96 Tissue Kit (Macherey Nagel, Hoerdt, France) and eluated in 100 µL of BE buffer. DNA concentration of these samples was measured using Quant-iT PicoGreen Kit (ThermoFisher, Waltham MASS, USA) according to manufacturer’s instructions.

Samples were then gathered with an equal DNA quantity into pools. DNA purification was performed for each pool with 1.8 times the sample volume in AMPure XP beads (Beckman Coulter, Fullerton CA, USA). Purified DNA were retrieved in 100 µL of ultrapure water. Pool concentrations were measured with Qubit using DNA HS Kit (Agilent, Santa Clara CA, USA). Final pool concentrations were as follow: 38.5 ng/µL for London Lab, 41.6 ng/µL for London Field, 40.3 ng/µL for German Lab and 38 ng/µL for Sweden Field.

Sequencing was performed using TruSeq Nano Kit (Illumina, San Diego CA, USA) to produce paired-end read of 2 × 150 bp length and a coverage of 25 X for London Lab, 32 X for London Field, 39.5 X for German Lab and 25.4 X for Sweden Field by Genotoul (Castanet-Tolosan, France).

### 2.4 Pool-seq data processing

The whole pipeline with the detail of parameters used is available on GitHub (https://github.com/chaberko-lbbe/clec-poolseq). Quality control analysis of reads obtained from each line was performed using FastQC (http://www.bioinformatics.babraham.ac.uk/projects/fastqc). The raw data have been submitted to the Sequence Read Archive (SRA) database of NCBI under BioProject PRJNA826750. Sequencing reads were filtered using Trimmomatic software v0.39 (Bolger et al., 2014), which removes adaptors. FastUniq v1.1 was then used to remove PCR duplicates (Xu et al., 2012). Reads were mapped on the *C. lectularius* reference genome (Clec_2.1 assembly, Harlan strain) performed as part of the i5K project (Poelchau et al., 2015), with an estimated size of 510.83 Mb. Mapping was performed using BWA mem v0.7.4 (Li and Durbin, 2009). Sam files were converted to bam format using samtools v1.9, and cleaned of unmapped reads (Li et al., 2009). The 1573 nuclear scaffolds were kept in this analysis, while the mitochondrial scaffold was not considered.

### 2.5 Detecting Single Nucleotide Polymorphisms

#### 2.5.1 Overall SNPs analysis

Bam files corresponding to the four populations were converted into mpileup format with samtools v1.9. The mpileup file was then converted to sync format by PoPoolation2 version 1201 (Kofler et al., 2011). 8.03 million (M) SNPs were detected on this sync file using R/poolfstat package v2.0.0 (Hivert et al., 2018) and the following parameters: coverage per pool between 10 and 50. In parallel, nucleotidic diversity (*π*) was computed using PoPoolation2 Variance-sliding.pl (default parameters).

Principal Component Analysis (PCA) was performed using R/pcadapt package v4.3.3 and “pool” type parameter (Privé et al., 2020). Fixation indexes (*F*_*ST*_) were computed with R/poolfstat for each pairwise population comparison of each SNP. Global SNP pool was then trimmed on minor allele frequency (*M AF*) of 0.2 (computed as *M AF* = 0.5 − |*p* − 0.5|, with *p* being the average frequency across all four populations). This relatively high MAF value was chosen in order to remove loci for which we have very limited power to detect any association with the resistance phenotype in the BayPass analysis. BayPass v2.3 (Olazcuaga et al., 2020) was performed with the standard covariate model, considering covariate data such as strain resistance statutes. The final dataset was thus reduced to 2.92M SNPs located on 990 scaffolds, and has been deposited on Dryad (doi.org/10.5061/dryad.9cnp5hqp6).

#### 2.5.2 Selecting outlier SNPs

Several filters were combined in order to detect SNPs potentially associated with insecticide resistance, hereafter called outlier SNPs. We first used R/poolfstat to identify the most differentiated alleles between the two most closely related populations, namely London Lab and London Field. Then, BayPass was used, as it takes into account the covariance of allelic frequencies between populations, arising because of common demographic history, and enabled to identify SNPs potentially under selection (i.e. SNPs showing extreme genetic differentiation). Thirdly, we selected alleles in a derived state compared to the allele carried by the reference Harlan strain, and in higher frequency in London Field than in London Lab. These steps are described more in detail below.

At first, to identify mutations potentially involved in resistance, we used fixation index (*F*_*ST*_) calculated with R/poolfstat for each pairwise population comparison for each SNP. SNPs with *F*_*ST*_ values belonging to the top 5% of the whole genome *F*_*ST*_ distribution computed between London Lab and London Field strains were considered as “outliers”. This comparison between London strains was chosen because of their high genetic proximity (see Results).

In a second step, we analyzed the whole dataset (the four strains) using the package BayPass. This package aims at identifying genetic markers associated with covariates (in our case, the resistance phenotype), while taking into account the shared history of populations. The contrast statistic C2 was computed to compare allele frequencies between resistant (London Field and Sweden Field) and susceptible strains (London Lab and German Lab). We corrected the p-values associated with C2 using R/qvalue version 2.26.0 to compute a local false discovery rate (lfdr) for each p-value. SNPs were first selected when having a lfdr below 0.2.

For each SNP across the four populations, the likelihood of the data was then estimated under a first model without selection, as well as under a second alternative model invoking selection. Bayes factor (BF) was computed to represent the ratio of these two likelihoods. A ratio equal to 2 indicates that the data favor the second model (with selection) twice as much as the first model (without selection). According to Kass and Raftery (1995), we considered SNPs having a log10(Bayes factor) > 0.5 as valuable candidates. In deciban (dB) units, used in BayPass (ie 10*log10(BF)), this corresponds to a threshold of 5.

Finally, we are assuming that the reference genome strain (Harlan strain), which is insecticide susceptible, carries the ancestral alleles for genes involved in insecticide resistance. Indeed, the Harlan strain was collected back in 1973 in the field, and thus has never been in contact with modern insecticides (Dang et al., 2017). From this perspective, we expected the frequency of alternative alleles (i.e. different from the reference genome allele at a position) to have increased over time under insecticide selective pressure. Therefore, only derived alleles with higher frequency in London Field were kept in the outlier list. SNPs that passed all combined filters described above were further considered as valid outliers, hence as associated with the resistance phenotype. A sliding window was computed for the number of outlier SNPs in 100 kb with a step of 10 kb, using sw function from the R package pegas v.1.1 (Paradis, 2010).

To locate SNPs in genes, we used the GFF3 file from NCBI (*C. lectularius* RefSeq genome, assembly accession GCF_000648675.2). It includes the type of feature and the coordinates of each genomic region (promoter, 3’ or 5’UTRs, intronic or exonic regions, and intergenic). Variants and their impact on protein sequences were analyzed using R/VariantAnnotation v1.34.0 (Obenchain et al., 2014) and R/GenomicFeatures v1.40.1 (Lawrence et al., 2013). Among the 2.92M SNPs, 1.52M (53%) were located within genes, with a huge majority inside intronic regions (1.45M, i.e. 51%). We identified 576 outlier SNPs (0.02%), 369 of which were located within genes (0.01% of all SNPs).

### 2.6 Mapping scaffolds on linkage groups

Based on a QTL RAD-seq analysis, Fountain et al. (2016) constructed a genetic map for *C. lectularius* assembly Clec_1.0. Nevertheless, the genome assembly of *C. lectularius* has been updated in the meantime (v1.0 to 2.1). In order to improve the genetic map produced, we decided to use Fountain’s RAD-seq data and R scripts to generate a genetic map based on the latest genome assembly. In order to assemble scaffolds into putative autosomes, hereafter called linkage groups (LG), we re-aligned Fountain’s RAD sequencing tags (available here http://dx.doi.org/10.5061/dryad.d4r50) with GSNAP version 2020-06-01 on the RefSeq genome version 2.1, rather than version 1.0 used by Fountain. By following the same pipeline, we were able to keep 338 out of 441 RAD markers, distributed over 14 linkage groups, as in Fountain analysis. Finally, out of 1573 nuclear scaffolds, 151 were kept, as containing uniquely mapped markers. The 14 putative autosomes obtained had an overall similar gene content as in Fountain et al. (2016) (Supp. fig. 4). While Fountain managed to map >65% of the genome on linkage groups, we only obtained 46% (i.e. 234.7 Mb out of 510 Mb). One explanation could reside in the higher fragmentation of the new genome version. Indeed, the latest version of the genome was split into more scaffolds (*Clec_1*.*0*: 1402 scaffolds, *Clec_2*.*1*: 1574 scaffolds see https://www.ncbi.nlm.nih.gov/genome/annotation_euk/Cimex_lectularius/101/). RAD markers therefore potentially fall into smaller scaffolds, which can lead to this loss of information.

An analysis of enrichment in outlier SNPs was performed on LG using R/stats default package (binomial test with Bonferroni correction for multiple testing). Then, to further identify smaller genomic regions of interest, enrichment tests were done at the scaffold level.

### 2.7 Functional analysis

In order to test whether some relevant biological functions were enriched in a set of candidate genes, we defined ten categories of genes potentially involved in resistance, similarly to what was done by Faucon et al. (2015), and benefiting from the extensive literature on the topic in several insects. Six of them corresponded to metabolic resistance: “Binding/Sequestration” (6 genes), “GST” (14), “CCE” (39), “UDPGT” (39), “ABC transporter/MRP” (55), and “P450” (56). The category “Other detox” (64) includes transcription factors, which could modulate detoxification enzyme expressions (Amezian et al., 2021). While “Cuticle” (113 genes) encompasses cuticular resistance genes, “Insecticide target and nervous system” (31) covers all the targets of the different insecticide families, as well as genes that could contribute to maintain neurological activity. Finally, bed bugs being hematophageous insects, they are subject to oxydative stress after a blood meal, and therefore adapted to deal with this stress. This feature could be advantageous for their defense against oxidative stress caused by pyrethroid exposure (Wang et al., 2016). We thus defined a “Redox homeostasis” category (14 genes). Based on gene annotation extracted from the GFF3 file (GCF_000648675.2), 431 out of the 13,208 genes (without *tRNA* and mitochondrial scaffold) were assigned to one of these categories (Supplementary table 1). Analysis of enrichment of candidate genes in each resistance category were performed by comparing their proportions in genomic regions of interest with whole genome, using R/stats default package (proportion test with FDR correction for multiple comparisons).

### 2.8 Structural variants

Emphasis was placed on two types of structural variants (SVs), duplications and inversions. London Field population was compared to the susceptible reference genome, and to the susceptible London Lab population, in order to detect SVs that could have been selected and underlie the resistance phenotype.

#### 2.8.1 Combining read depth and pairs orientation to detect SVs

The presence of SVs was inferred based on abnormal read pair orientation and/or distance (insert size), and read depth variation. We used poolCNVcomp (Schrider et al., 2013, 2016; North et al., 2020), a series of scripts designed for the detection of tandem duplications in pool-seq data, and adapted them to detect inverted duplications and simple inversions as well. The first step identifies read pairs having abnormal orientation and extreme insert size (top 1% distribution) in each population. In a tandem duplication, read pairs that are falling between the two copies will both map on the reference genome in a reversed way (i.e., in opposite direction). For inverted duplication, one of the two paired reads will be in the wrong orientation. The 2nd step groups them together if they fall inside the same genomic region, using parameter insertSizeDiffCutoff (standard deviation of insert sizes distribution respectively to each scaffold, multiplied by four). At the end of these two first steps, candidate structural variants have been delineated, in comparison to the reference genome. The 3rd step then matched corresponding structural variants between population pools, using the parameter distancecutoff (redefined as the sum of insertSizeDiffCutoff from populations A and B), to determine if the coordinates are close enough to represent the same event in resistant and susceptible populations.

The 4th step aims at computing read depth in each pool. During this process, repeated regions were masked. To obtain those repeated regions, we performed a blast of transposable elements detected in *C*.*lectularius* by Petersen et al. (2019) on the latest genome assembly, i.e. *Clec_2*.*1*. All hits with more than 80% nucleotidic identity were considered as repeated regions and masked.

Next, in North et al. (2020) 4pt5 step, we used the coverage data calculated for each pool to test whether London Field had more copies compared to London Lab in putative duplications. Under this hypothesis, the coverage ratio London Field/London Lab on a locus of interest should be disproportionately high compared to a null expectation. To determine a threshold, we estimated empirically the read depth ratio London Field/London Lab under the null hypothesis of no amplification between London Field and London Lab strains. This expectation was computed for a set of windows of different sizes (500 bp to 2.5 Mb, n=10 000), randomly sampled in the genomes. We then compared observed values with the corresponding null distribution (for a given size class), and considered that events with ratios falling above the 75% upper quantile were valid candidates for high copy number amplifications.

We finally filtered out events smaller than the size class of 3.5 kb (half the median size of genes, being 7 kb), leading to our global set of events. We used the same script to identify inversions not associated with duplications. The single difference with the previous pipeline was that the coverage ratio had to be between the 25% and 75% quantiles.

The presence of transposable elements may interfere with the calling of SVs due to multiple mapping. We therefore removed structural variants if their coordinates on extremities were overapping for more than 100 bp with TE (on 150 pb).

#### 2.8.2 Identifying highly differentiated SVs

A rough estimate of the allelic frequency of each event was done by computing the coverage in the 150 bp inward flanking regions of each event for each strain, using samtools v1.9. For each pool, we then estimated the frequency of reads supporting the SV by dividing the number of reads with peculiar orientation by the mean value of the coverage in the flanking regions. Events showing a higher frequency in London Field than in London Lab were selected.

We then calculated the *F*_*ST*_ index of differentiation based on these estimated frequencies. *F*_*ST*_ falling in the top 10% were considered as valid candidates (*q*_90_ = 0.040 for inversions, 0.040 for inverted duplications and 0.036 for tandem duplications).

We tested whether scaffolds were enriched with structural variants or not, by comparing the observed number of SV per scaffold with the expected one under the hypothesis of homogeneous distribution among the scaffolds (expected number being the total number of SV divided by the total size of scaffolds, multiplied by each scaffold size). Scaffolds having less than 5 expected events were then removed. We then computed a *χ* ^2^ test for each scaffold (1 d.f.), and corrected the associated p-values using Bonferroni method. The same method was used to test whether scaffolds were enriched in a specific type of structural variant (tandem duplications, inverted duplications, or inversions).

Finally, we detected for each category events that were overlapping with genes. To do so, we extracted all genes falling into windows encompassing an event and 1 kb further upstream and downstream. We also looked at SVs affecting candidate resistance genes.

## 3 RESULTS

### 3.1 Resistance phenotype of the four strains

We tested the resistance statutes of the four *C. lectularius* strains to the pyrethroid insecticide deltamethrin. As expected, the two recently collected field strains showed higher deltamethrin resistance as compared to the two ancient lab strains (Figure 1.A).

**FIGURE 1.**
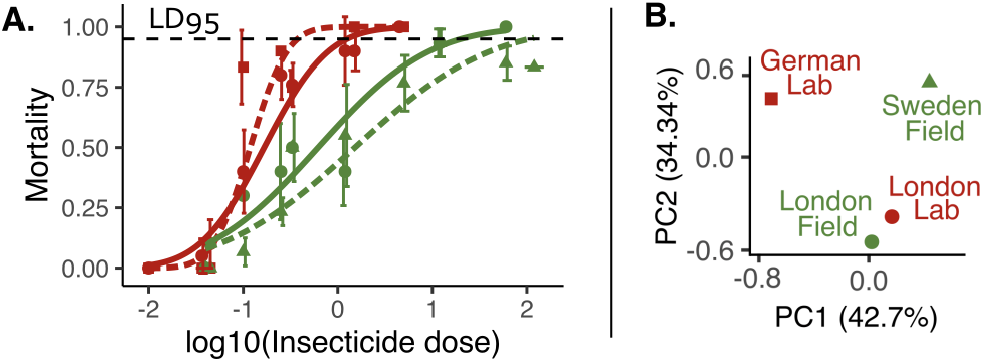
Analysis of the four bed bug strains used (German Lab: red square, London Lab: red round, London Field: green round, Sweden Field: green triangle). A. Dose-response curves to the insecticide deltamethrin exposure in bed bug populations German Lab (dashed red line), London Lab (red line), London Field (green line) and Sweden Field (dashed green line). Average mortality percentages are shown for each log10-transformed insecticide dose (in ng/µL) with standard deviation across replicates. Lines correspond to probit transformations of the data. B. Projection of four bed bug populations on the top two principal components using PCA on 8.03 million SNPs.

German Lab and London Lab had the lowest LD_95_ values, respectively 0.4 ng and 1.3 ng of deltamethrin per bed bug. On the contrary, London Field and Sweden Field had higher LD_95_, respectively 21.0 ng and 69.2 ng (Supplementary Table 2). Resistance ratios to this dose (RR_95_) between field and lab strains were all >10 X (Table 1), which is usually considered as high resistance ratio (Mazzarri and Georghiou, 1995).

Comparisons revealed that all strains had significantly different phenotypes in terms of insecticide resistance, exception made of Sweden Field versus London Field and London Lab. Overall, these results confirmed the resistance statutes of the two field strains (R) compared to the two susceptible lab strains (S).

### 3.2 Genome-wide genetic differentiation

8.03 million (M) SNPs, distributed over 1,077 scaffolds, were detected. The genetic diversity, measured as *π*, was very similar across the four populations (Table 2). After PCA, the four populations were projected onto two principal components, PC1 and PC2, explaining respectively 42.71% and 34.34% of variance (Figure 1.B). The four strains clustered according to their geographic origin, rather than their age or their deltamethrin resistance statutes. The historical and modern strains collected in London were noticed to be very close to each other on the 2D plot, whereas strains collected in different countries were distant (respectively Germany, Sweden and United Kingdom). This observation suggests that London Lab and London Field shared a common genetic background comparatively to the other strains. Average fixation indices (*F*_*ST*_) were then computed to provide a measure of pairwise population genetic differentiation. As expected from the PCA analysis (Figure 1.B), the mean *F*_*ST*_ calculated between London Field and London Lab was the lowest (*F*_*ST*_ = 0.018, i.e. 1.8%), compared to the other comparisons which revealed much higher levels of differentiation (between 4.8% and 8.9%, Table 2). It therefore confirmed the weak genetic differentiation between the two strains from London. Both PCA and *F*_*ST*_ analysis thus indicate that the comparison between London Lab and London Field is particularly relevant to identify candidates potentially involved in insecticide resistance.

**TABLE 1.** Pearson’s chi square goodness-of-fit (pgof) statistical test on insecticide deltamethrin LD_95_ for *Cimex lectularius* population comparison, with associated standard errors and resistance ratios. Significance levels for statistical tests are indicated, based on p-values (not significant (-), <0.05 (*), <0.001 (**)).

| Population comparison | Statistical pgof test | Standard error | Resistance ratio |
| --- | --- | --- | --- |
| London Lab - London Field | 2.95 (*) | 0.414 | 17 X |
| German Lab - London Field | 4.34 (**) | 0.411 | 60 X |
| London Lab - Sweden Field | 1.48 (-) | 1.17 | 55 X |
| German Lab - Sweden Field | 1.96 (*) | 1.17 | 199 X |
| German Lab - London Lab | 6.81 (**) | 0.082 | 4 X |
| London Field - Sweden Field | 0.42 (-) | 1.24 | 3 X |

**TABLE 2.**
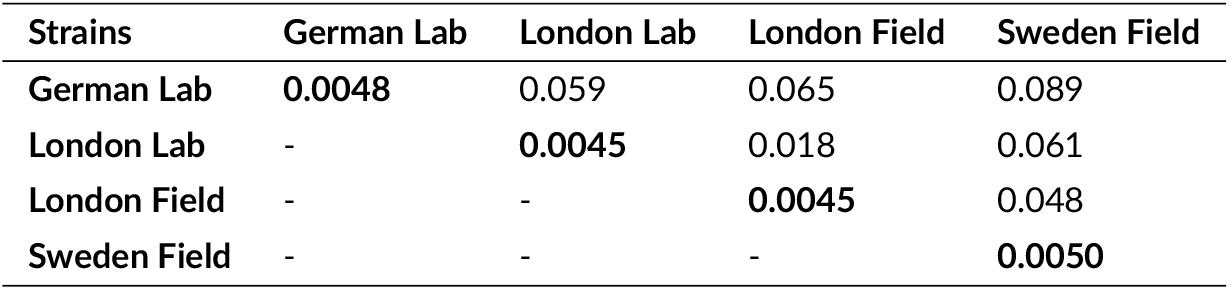
Mean *F*_*ST*_ values for pairwise comparisons of *Cimex lectularius* strains, and within-strain nucleotidic diversity (*π*, in bold).

### 3.3 Overview of SNPs

Within the 2.92M SNPs that passed the QC filters, 576 outlier SNPs were identified (using top 5% *F*_*ST*_ *LL* −*LF*, BayPass metrics and *F*_*al t*_ *LF* > *F*_*al t*_ *LL*, see Methods for detail). Among them, 369 were detected within genes. With only 7 outliers in putative resistance genes, there was no significant overall enrichment. Only one non-synonymous outlier SNP was found out of 10,318 non-synonymous SNPs. This non-synonymous mutation was detected in a homeobox protein one cut (LOC106666929) located in a scaffold not included in any linkage groups and not particularly enriched for outlier SNPs (NW_019392707.1, Supplementary figure 3).

#### 3.3.1 Distribution of outlier SNPs

Among the 576 outlier SNPs, 261 were located within linkage groups (LGs). To identify LG of particular interest, we first tested whether some were enriched with outlier SNPs, by comparing the proportion of outlier SNPs for each LG to the average value computed for the entire genome.

LG 8 and 11 were significantly enriched with outlier SNPs (indicated by a star next to LG number in fig. 2). In LG 8, despite a global 2.18 X enrichment, no scaffold was significantly enriched for outlier SNPs, nor were any outliers detected in putative resistance genes.

**FIGURE 2.**
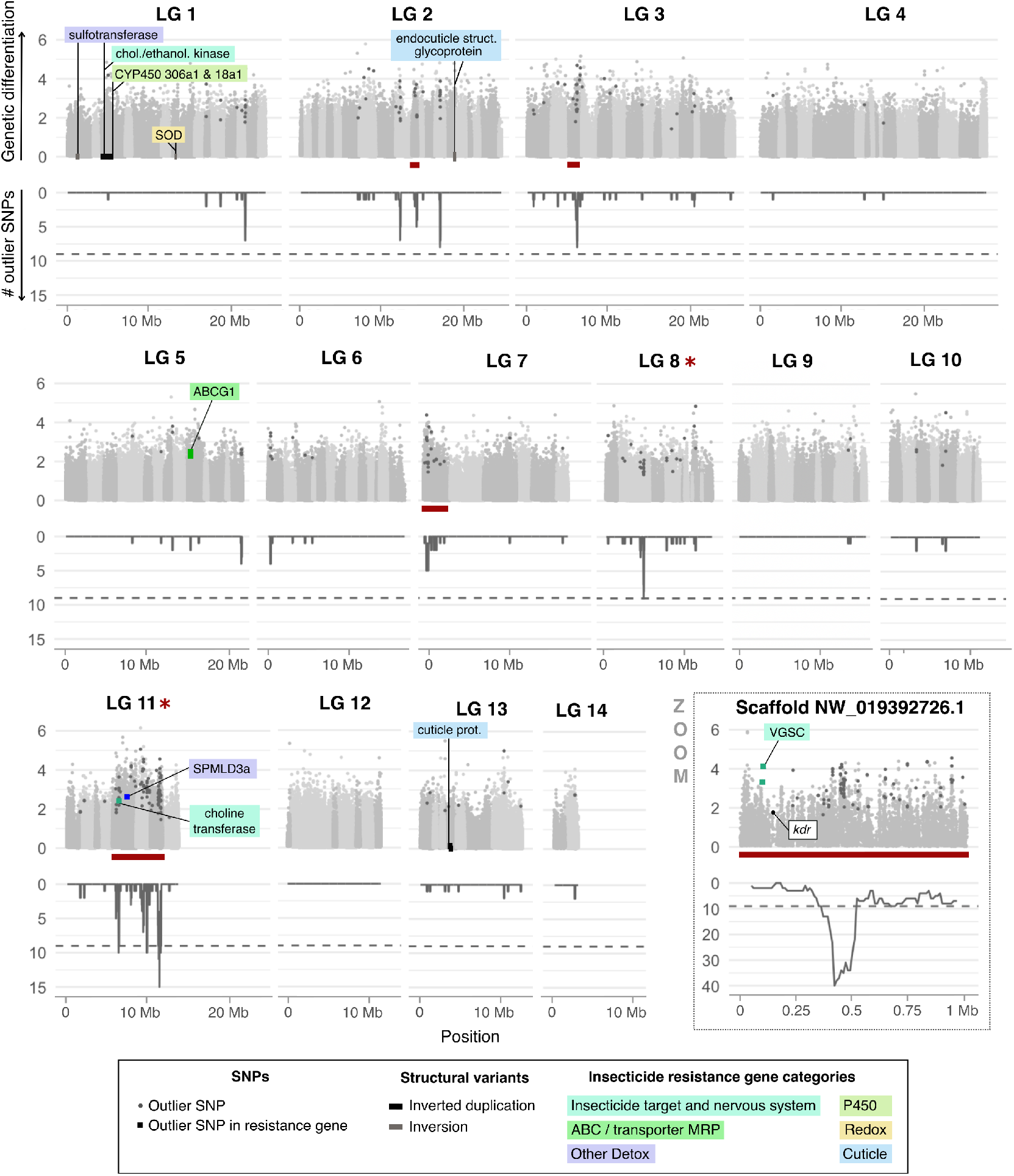
Outlier SNPs in *Cimex lectularius* linkage groups. Each plot represents a linkage group (LG), exception made of scaffold NW_019392726.1 (unplaced, zoomed on). Within a LG, scaffolds are delimited by different shades of grey. In the upper part of each plot, genetic differentiation is represented with empirical p-values, computed on *F*_*ST*_ index between London Lab and London Field populations for each SNP in -log10 scale. Outlier SNPs (see Materials & methods) are distinguished by a darker grey. Square-shaped points are the outlier SNPs that fall within candidate resistance genes and are colored according to those genes’ categories. Their protein products are labeled. Although not an outlier SNP, the location of the non-synonymous *kdr* L925I mutation was given as an indication. In the lower part, the number of outlier SNPs counted in each 100 kb sliding window with a step of 10 kb is represented. Dashed line stands for a threshold of 2 outlier SNPs, which delineates quantile 99%. The positions of structural variants (inverted duplications in black and inversions in grey) overlapping with resistance genes and with top 10% *F*_*ST*_ values are displayed on the *x*-axis (y=0). Annotations of overlapping resistance genes are labeled and coloured according to the resistance category.

Still, a large number of outliers were found on the second scaffold, namely NW_019393011.1, mainly inside two uncharacterized genes (106663904 and 106663882) and a small conductance calcium-activated potassium channel protein (106663893). This last gene was not included in our pre-defined resistance genes categories, but can still be involved in insecticide resistance through its potential role in synaptic functioning (Vergara et al., 1998).

By far, the highest enrichment was observed for LG 11 with a 4.86 X enrichment (p.value = 1.66e-34, Supplementary figure 1). Interestingly, most of the outlier SNPs clustered within three consecutive scaffolds within this LG (NW_019392721.1, NW_019392763.1 and NW_019942502.1.1). These three scaffolds were also significantly enriched for outlier SNPs (underlined in red in fig. 2). Within this approximately 6 Mb region, we found two outlier SNPs in the intronic regions of two putative insecticide resistance genes, both on scaffold NW_019392721.1: a choline O-acetyltransferase (106664013), an enzyme that catalyzes the biosynthesis of acetylcholine, and an acid sphingomyelinase-like phosophodiesterase 3a (SMPDL3a, 106664028), which may be involved in insecticide detoxification (Zhang et al., 2012). However, many other candidate resistance genes were found inside this region, thus being further explored below (see Section 3.3.2).

Other scaffolds in LG detected as enriched were rather dispersed, without close proximity to putative resistance genes. Finally, two outlier SNPs were detected in an ABC transporter G, located on LG 5 which is not particularly enriched in outliers, neither at the scale of the whole LG nor at the scale of its scaffolds. ABC transporter G may be involved in the excretion of insecticides out of the cells (Dermauw and Van Leeuwen, 2014).

Eight scaffolds not included in the 14 linkage groups were significantly enriched for outlier SNPs (Supplementary figure 3). Two of these scaffolds carried outlier SNPs in putative resistance genes. A first outlier SNP was detected in an intronic region of a beta-14-galactosyltransferase 4-like on scaffold NW_019392678.1. Galactosyltransferase is a type of glycosyltransferase, an enzyme capable of contributing to xenobiotic detoxification through the conjugation of an electrophilic group to xenobiotics or to their metabolites (Nagare et al., 2021).

Two intronic outlier SNPs were also detected in the voltage-gated sodium channel gene (*VGSC*, 106667833), on scaffold NW_019392726.1, in close proximity with the *kdr* mutation. The *kdr* non-synonymous SNP of the *VGSC* L925I (leucine to isoleucine in position 925, i.e. position 148,668 on NW_019392726.1), was indeed further identified. Frequencies of the derived allele were as follows: 0% in London Lab, 11.1% in German Lab, 44.4% in London Field, and 50% in Sweden Field. This locus was part of the top 1.6% *F*_*ST*_ in the London Lab *versus* London Field comparison. The *kdr* mutation has been extensively studied in insects, including bed bugs, and is associated with pyrethroid resistance (see review Dang et al., 2017). Although our analysis did not consider *kdr* mutation as an outlier *per se, kdr* mutation still belongs to a peculiar 1 Mb genomic region (i.e. the scaffold size) enriched for outlier SNPs.

#### 3.3.2 Detection of a superlocus enriched in both outlier SNPs and resistance genes

Three consecutive scaffolds enriched for outliers were detected on the LG 11 (NW_019392721.1, NW_019392763.1 and NW_019942502.1, Figure 3). In the following, this 6 Mb genomic region is referred to as a “superlocus”. Among the 138 genes located within this superlocus, nine were part of a resistance gene category, which represents a significant enrichment for resistance genes when compared to the entire genome (prop.test, p-value = 0.027). More precisely, two categories of resistance genes were significantly enriched: *GSTs* (corrected p-value = 3.0e-9, 3 genes in the superlocus out of 14 in the whole genome), and *P450s* (corrected p-value = 0.029, 3 genes in the superlocus out of 56 in the whole genome). The same trend was observed for the category ‘Insecticide target and nervous system’, although this was only marginally significant (corrected p-value = 0.063, 2 genes out of 31). This result suggests a clustering of genes potentially involved in insecticide resistance in this superlocus.

**FIGURE 3.**
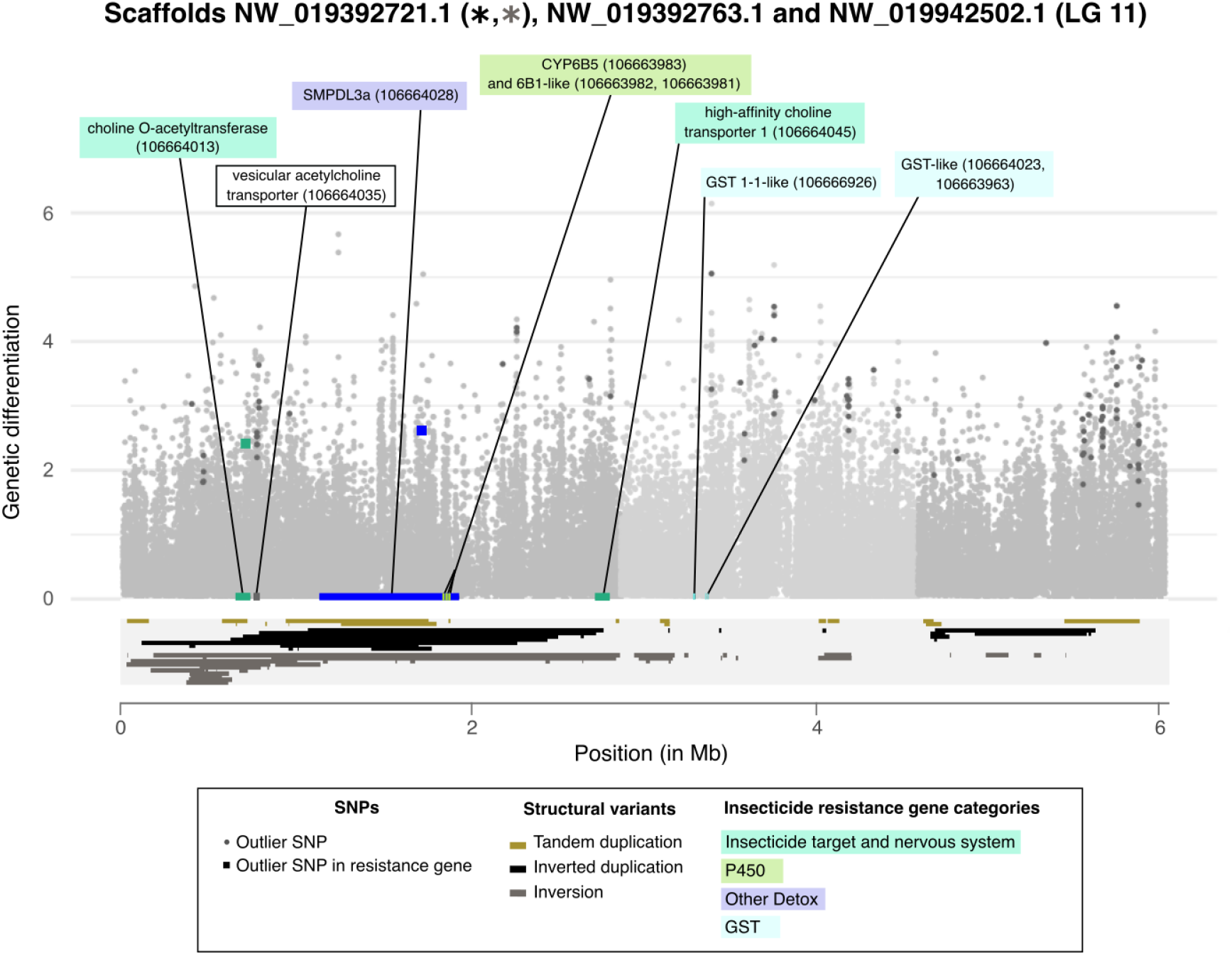
Architecture of *Cimex lectularius* superlocus on LG 11. Within this superlocus, scaffolds NW_019392721.1, NW_019392763.1 and NW_019942502.1.1 are delimited by different shades of light grey. Genetic differentiation is represented with empirical p-values, computed on *F*_*ST*_ index between London Lab and London Field populations for each SNP in -log10 scale, depending on their position within each scaffold. Outlier SNPs (see Materials & methods) are distinguished by a darker grey. Square-shaped points are the outlier SNPs that fall within resistance genes, and are colored according to those genes’ categories. Resistance gene positions are indicated on the *x*-axis and colored accordingly to their categories. Protein product of resistance genes are labeled, along with the NCBI accession number of the genes between brackets. One vesicular acetylcholine transporter, carrying outlier SNPs and in close proximity to choline-O-acetyltransferase, was also labelled likewise. The positions of structural variants were represented inside a light grey box under the *x*-axis. Inverted duplications are in black, inversions in grey, and tandem duplications in gold. A star of the same color was added next to the scaffold name if it was enriched in a type of event.

Indeed, the three *P450s* detected in the superlocus were located next to each other, all belonging to the sub-family *CYP6B* (106663981, 106663982 and 106663983). Members of the CYP6B P450 sub-family have been shown to metabolize a wide range of natural and synthetic xenobiotics in insects, including pyrethroids, as in *Helicoverpa armigera* (Shi et al., 2021). The same pattern was observed for the three *GST* detected, clustered on scaffold NW_019392763.1 (106666926, 106664023 and 106663963). *GST* have also been frequently associated with pyrethroid resistance, as for *Anopheles sinensis* (Tao et al., 2022).

Interestingly, despite none of these genes carried outlier SNP, they showed 23 non-synonymous SNPs with an alternative allele frequency higher in London Field than in London Lab (Supplementary table 3). Four of these non-synonymous SNPs were also part of the top 5% *F*_*ST*_, with two for each P450s gene 106663983 and 106663982.

Because the abundance of non-synonymous mutations may be the consequence of adaptive evolution, we calculated the ratio 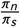 all the genes of the genome. The 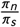 for the three *CYP6B* genes were in the top 5% of the distribution (quantile 95% = 0.88, mean = 0.19, versus 1.07, 1.25 and 1.50 respectively for genes 106663981, 106663982 and 106663983). This result supports that adaptive evolution has shaped the polymorphism for these P450s.

### 3.4 Structural variants associated with resistance

Structural variants (SVs) may play an important role in insecticide resistance (Assogba et al., 2015). We detected 2,497 SVs with higher frequency in the resistant strain London Field compared to the susceptible London Lab, encompassing 1,338 inversions, 650 inverted duplications, and 509 tandem duplications.

Of the 2,497 SVs detected, 143 were overlapping with 130 putative resistance genes (out of 431 in the whole genome), which is more than what is expected by chance (p-value=3.53e-11). This indicates an enrichment of SVs in genes possibly related to insecticide resistance. Among SVs overlapping with candidate resistance genes, 80 genes were found within inversions, 50 in inverted duplications and 30 in tandem duplications.

Seven scaffolds were significantly enriched in SVs compared to what was expected based on their respective lengths, if one assumes that events were equally distributed among scaffolds: NW_019392673.1 (LG 1), NW_019392754.1 (LG 7), NW_019392785.1 (LG 10), NW_019392721.1 (LG 11), NW_019392782.1 (LG 12), NW_019392665.1 (LG 13) and NW_019392786.1 (unplaced) (details in Supplementary figure 2). Among them, the only scaffold that was significantly enriched both for outlier SNPs and SVs was the scaffold NW_019392721.1 (Supplementary figure 2), located within the superlocus of LG 11 and encoding the three *CYP6B* with unusual high 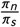 ratios (Figure 3).

For each SV category (inversion, inverted duplication, tandem duplication), events falling above the 90% quantile of *F*_*ST*_ were retained, leading to 227 SVs overlapping with 281 different genes. Among them, ten SVs overlapped with 18 resistance genes. Seven cuticular genes were identified (in three inversions and one inverted duplication), showing an enrichment in this resistance category (p-value = 0.01). The largest event detected was a 1.5 Mb inverted duplication located on LG 1 (NW_019392679.1). This event overlapped with four putative resistance genes: a sulfotransferase, a choline kinase, and two P450s (Supplementary table 4). The five other events overlapped genes possibly involved in redox homeostasis (SOD and NADPH-dependent oxidoreductase), in metabolic resistance (UDPGT on scaffold NW_019392656.1) and in other detox (sulfotransferase).

## 4 DISCUSSION

Insecticide resistance evolution seems to be the key factor in the re-emergence of the common bed bug *C. lectularius* (Davies et al., 2012). By using targeted sequencing (Zhu et al., 2010), transcriptomic analysis (Zhu et al., 2012; Mamidala et al., 2012; Koganemaru et al., 2013), or QTL mapping (Fountain et al., 2016), several loci have been associated with insecticide resistance (reviewed in Dang et al. 2017). In this paper, we provide a genome-wide analysis to identify underlying sequences and additional regions of the genome possibly involved in insecticide resistance. By comparing the whole genomes of two susceptible and two resistant lines using pool-seq data, we identified outlier SNPs and structural variants (inversion, tandem duplications, and inverted duplications) highly differentiated in the resistance phenotype, in close proximity with candidate resistance genes. In addition to providing novel insecticide resistance data, the genomic characterization of CimexStore strains will hopefully benefit future customer-researchers that regularly use these strains for their research.

### 4.1 Relatively low pyrethroid resistance level

Using topical application assays, we quantified the deltamethrin resistance level of the four strains. As expected from the collection dates, the two ancient strains (LL and GL) were highly susceptible to deltamethrin. In most studies, the Harlan strain is used as a insecticide susceptible reference strain (Dang et al., 2017). The susceptibility of the LL and GL strains was comparable to that of the Harlan strain (0.34 ng and 0.15 ng per insect respectively for London Lab and German Lab, compared to 0.15 ng for the Harlan strain (Cáceres et al., 2019).

This high sensitivity is expected since these two “lab” strains were collected before the re-emergence of bed bugs, as was the Harlan strain. Conversely, the two relatively-recent strains (LF and SF), respectively collected in 2008 and 2015 were much more resistant to deltamethrin. However, their resistance ratios were not particularly high (17 to 199 fold based on LD_95_), as compared to Cáceres et al. (2019), where field strains from Barracas and Pergamino were respectively 6,945 and 10,178 times higher than their susceptible reference. Resistance ratio should be compared with caution, since the bioassays or analytical methods can be widely different and have an impact. However, very high RR phenomena have frequently been observed for recently collected field strains (see review Dang et al. 2017), but often decline after rearing in the laboratory without insecticide exposure (Polanco et al., 2011). The existence of an evolutionary cost of resistance in the absence of insecticide exposure has been frequently described in insects, and may explain the decrease in frequency of resistance alleles since the establishment of the strains (Kliot and Ghanim, 2012). This phenomenon may explain the relatively low level of resistance for the two CimexStore “field” strains purchased, for which no information was provided regarding insecticide selection pressure during the breeding in laboratory conditions.

### 4.2 Non-synonymous outlier SNP in a transcription factor

Adaptation of organisms to their environment may rely on protein evolution thanks to non-synonymous mutations, and/or thanks to expression changes involving mutations in regulatory sequences (Wang et al., 2005). Among the 576 outlier SNPs, only one was associated with an amino-acid change. The encoded protein, annotated as an homeobox protein onecut (106666929 on Supplementary fig. 3), is composed of a onecut domain and one homeodomain which together confer DNA binding properties (Bürglin and Affolter, 2016). Homeobox proteins onecut are transcription factors involved in many pathways including nervous system differentiation (Nguyen et al., 2000). Several transcription factors have been associated with resistance to insecticide, as in *Plutella xylostella* (Zhao et al., 2018). Yet, how this particular transcription factor could be involved in insecticide resistance remains unclear.

### 4.3 A superlocus involved in insecticide resistance

The most striking result obtained from this genome-wide analysis is the identification of a 6 Mb genomic region, showing an unusual high genetic differentiation and associated with the resistance phenotype of the four strains (BayPass approach). Several phenomena may explain such local genetic differentiation. Large genetic differentiation islands may evolve because of a local reduction in recombination rate due to the effect of linked selection (Burri et al., 2015). However, if this factor was at play, then we would expect a higher genetic differentiation in this genomic region for all comparisons, including between the two “field” strains or between the two “lab” strains. This was not the case, since this genomic region was particularly differentiated in the London “Field” versus London “Lab” comparison, but not in other comparisons (Suppl. fig. 5). High local genetic differentiation may thus be the consequence of differential selection. In this case, the selective factor is most likely the exposition to insecticides that recent field strains experienced, as compared to older “lab” ones. This region was part of LG 11 and was composed of three scaffolds. Those scaffolds were enriched with outlier SNPs, and one of them was also enriched with structural variants.

The detection of this superlocus is consistent with a previous study from Fountain et al. (2016) who constructed a linkage map and performed a QTL mapping for pyrethroid resistance in *C. lectularius*. Their QTL analysis based on RAD tag sequencing identified a single peak associated with resistance. Due to the relatively low number of markers (n=334), to the relative small number of recombinants screened (n=71) and to the small number of generations allowing recombination (F2), the resolution achieved was relatively low and led to the identification of a large 10.8 Mb region. This regions was located on their LG 12 and contains the highly expected insecticide resistance candidate *VGSC*, as well as many other genes, including putative resistance genes (P450 and GSTs). This large locus explained most of the variance in insecticide resistance in their experimental display. Comparing the two LG belonging to the two linkage maps indicated that the majority of the genes found on our-LG 11 was shared with Fountain-LG 12 (Supplementary figure 4). However, the scaffold carrying the *VGSC* gene which was part of Fountain-LG 12 was not included in our-LG 11. Instead, the scaffold was left unplaced in any LG. The explanation for this discrepancy is simple. In the previous genome assembly Clec_1.0, the *VGSC* gene belonged to a 21 Mb scaffold (NW_014465016.1) containing 6 RAD tags (See Supplementary table S2 of Fountain et al. 2016), that allowed Fountain et al. (2016) to incorporate *VGSC* gene into their LG 12. This scaffold was then split in smaller pieces in Clec_2.1 assembly, leaving *VGSC* in a 1 Mb-scaffold not carrying any RAD marker anymore (NW_019392726.1). This difference in assemblies thus prevented its integration into the new genetic map.

The second version of the assembly was obtained by using Redundans on top of the first assembly (Thomas et al., 2020). Redundans (Pryszcz and Gabaldón, 2016) is designed for solving problems linked to the presence of heterozygosity in genome sequencing projects. Based on read depth and paired-end information, it identifies putative “heterozygous contigs” which correspond to different allelic version of a single locus. They are then merged in order to produce as far as possible a homozygous version of the genome. Although the reason explaining the split of the initial 21 Mb scaffold (NW_014465016.1) into smaller pieces is unclear, we may hypothesize that some of the contigs that composed this scaffold were detected as “heterozygous contigs” ultimately leading to the observed split. Future investigations should certainly clarify this point and more generally clarify the structure of the haplotypes in this genomic region.

We note however that in the 21 Mb scaffold NW_014465016.1 of Fountain-LG 12, *VGSC* was flanked by two genes that are both included in our LG 11 (a carboxyl terminal-hydrolase on scaffold NW_019942502.1, and a glutathione S-transferase on scaffold NW_019392763.1). The position of the scaffold containing *VGSC* gene (NW_019392726.1) may thus possibly be between the two scaffolds previously mentioned, in the middle of the superlocus. If this is the case, it would further reinforce the strong signal of genetic differentiation in this genomic area.

Independently of its exact genomic localization, we expected the mutant sites of the *VGSC* gene (with their socalled knockdown resistance -*kdr*alleles) to be part of our outlier list. This was expected since *kdr* mutations are responsible for or are associated with resistance to pyrethroids in various arthropod pests (Dong et al., 2014), including bed bugs (Dang et al., 2017). Additionally, CimexStore website indicates the presence of the common *kdr* mutant L925I in the two field strains. Accordingly, we detected this mutation in our dataset. However, this particular site was not considered as an outlier SNP in our analysis. This site fulfilled only two out of three criterion used since (i) the *F*_*ST*_ associated with this site was part of the top 5% in the London strains comparison and (iii) the *kdr* mutant was derived compared to the reference genome, but (ii) it was filtered out by the BayPass analysis, indicating an overall weak association with resistance phenotype in the four strains. Several scenarios may explain this unexpected result. First of all, we can hypothesize that this mutation is not as crucial for insecticide resistance in *C. lectularius* as previously thought. Under this scenario, other mechanisms such as detoxification, cuticle thickening or behavioural avoidance would be more central in bed bug insecticide resistance. Importantly however, since these strains have been reared in the lab since years by the private company CimexStore, we do not exactly know under which conditions they have been maintained. One crucial point to keep in mind is that *kdr* mutants are highly counter-selected in the absence of insecticide (Freeman et al., 2021; Wang et al., 2021) and may thus decrease in frequency if not regularly exposed to insecticides. We can therefore question whether the frequency estimated here reflects the true initial frequency of that mutation in the different strains. If present, this effect may thus reduce the genetic differentiation observed on this locus (and surrounding region) between field “resistant” and ancient lab “susceptible” strains. Despite the fact that our analysis did not consider *kdr* mutant as part of the outlier list, we found two outlier SNPs in an intron of the *VGSC* gene. Additionally, *VGSC* was located next to a region containing the highest density of outlier SNPs, with up to 40 outlier SNPs per 100 kb window (Figure 2). Altogether, these last observations indicate that *VGSC* belongs, at least, to a peculiar 1 Mb region in relation to insecticide resistance.

### 4.4 A peculiar cytochrome P450 cluster

A cluster of three P450 genes (two *CYP6B1*-like and one *CYP6B5*-like) was detected in scaffold NW_019392721.1, in the middle of the LG11 superlocus. No outlier SNP were detected in any of these loci. However, four SNPs were part of the top 5% *F*_*ST*_ distribution for the London strains comparison. In addition, this cluster was characterized by an unexpectedly high level of non-synonymous mutations (n=19). Accordingly, the 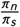 ratio for all three genes was in the top 5% or the genome-wide distibution, suggesting that selection may have acted on these loci in a recent past. Similar increased rate of evolution has been observed for *CYP6B56* of the navel orangeworm *Amyelois transitella* (Calla et al., 2021). In this species, *CYP6B56* is likely involved in the pyrethroid bifenthrin resistance, since elevated expressions of *CYP6B56* were observed after insecticide exposure in both resistant and susceptible populations (Calla et al., 2021).

### 4.5 Structural variants might contribute to shape resistance phenotype

Structural variants may contribute to adaptation, notably through inversions or duplications. These structural changes can affect the coding sequences as well as the expression levels of the genes, thus playing a crucial role in evolution and adaptation (North et al., 2020). CNVs in particular may impact the level of expression of the genes through a dosage mechanism (Kondrashov, 2012). In addition, CNVs can allow alternative variants of a gene to appear in tandem, through gene duplication. This may be selected for, especially when the allele encoding resistance also carry a fitness cost in the absence of insecticide. In that case, the second nearby copy may compensate for this cost and “heterogeneous” tandem duplications may be selected for (Labbé et al., 2007; Assogba et al., 2015). Despite their potential importance, SVs have received less attention than SNPs, likely because they are harder to identify. Population-level genome-wide analyses of SVs are thus rare, and the extent of their impact on evolution remain overlooked (Lucas et al., 2019). However, some SVs have been associated with insecticide resistance in insects, such as in *Drosophila* (Schmidt et al., 2010), in several mosquito species (Faucon et al., 2015; Weetman et al., 2018; Lucas et al., 2019; Cattel et al., 2020, 2021), or in the fall armyworm with the identification of a CYP cluster (Gimenez et al., 2020). To the best of our knowledge, the present study is the first to investigate SVs in an agnostic genome-wide fashion in bed bugs. We found that SVs are enriched in putative resistance genes. A similar test of enrichment focusing on copy number variants (CNVs) has been conducted on the mosquito *Anopheles gambiae*, using the Ag1000G Phase 2 dataset (1142 whole genome sequences of wild-caught mosquitoes collected from 16 populations) (Lucas et al., 2019). The authors found an enrichment for CNVs in gene-containing regions. More specifically, they found an enrichment in putative metabolic resistance gene families. Similar result has been observed in *Drosophila* (Schrider et al., 2016) and *Aedes aegypti* (Faucon et al., 2015; Cattel et al., 2020, 2021). Additionally, several independent amplification events were frequently found as covering the same genes, possibly segregating within the same populations (Lucas et al., 2019). We reached the same conclusion, with some genomic locations highly enriched with overlapping events. This was in particular the case for scaffold NW_019392721.1 in LG 11 which contained several candidate genes, including a cluster of three P450, and that was enriched both with outlier SNPs and SVs. Furthermore, we found a few set of events overlapping resistance genes with high *F*_*ST*_ values (top 10%, max: top 1%, see Supp. table 4). Among them, a significant enrichment for cuticle genes was detected. Although the functional importance of these SVs in the resistance phenotype remains to be further investigated, these results support their potential role in adaptation to insecticides in bed bugs.

### 4.6 A supergene involved in insecticide resistance?

A significant enrichment for genes potentially involved in insecticide resistance was observed in the 6 Mb superlocus. Colocalization of genes involved in detoxification in clusters is frequently observed in insects, including the mosquito *Anopheles gambiae* (Ranson et al., 2002). This clustering is probably the consequence of local duplications or of translocation of genes into the cluster. The discovery of a single large (at least 6 Mb long) locus probably involved in insecticide resistance is uncommon in the literature. In most cases reported to date, the loci identified by population genomics analysis are relatively small (<200 kb in Weedall et al. 2019) and numerous (Pélissié et al., 2022), and no evidence of genomic clustering among those loci is provided. However, few exceptions exist. In the navel orangeworm (*Amyelois transitella*), an exceptionally large region with low polymorphism, extending to up to 1.3 Mb was detected in a DDT and bifenthrin resistant population (Calla et al., 2021). This region contained several genes likely involved in insecticide resistance, such as VGSC and a cytochrome P450 gene cluster. The genomic co-localization of genes that are selected for in the same environment (i.e. under insecticide exposition) may theoretically lead to the emergence of so-called “supergenes” if a subsequent reduction of recombination between beneficial alleles occurs (Schwander et al., 2014). In a spatially or temporally heterogeneous environment (in that case treated versus untreated households), co-transmission of alleles may be selected for. Several fascinating examples of such evolution have been reported, from the classical case of Butterfly mimicry in the genus *Heliconius* (Le Poul et al., 2014) to the evolution of prairie or forest ecotypes of deer mice *Peromyscus maniculatus* (Hager et al., 2022). Although our analysis does not provide any evidence of a reduction in recombination in this superlocus (Supp. fig. 6), it is intriguing to observe such enrichment for putative resistance genes in a highly differentiated and large genomic island. Additionally, our dataset suggests an enrichment for structural variants, especially inversions, in one of the three scaffolds composing this superlocus (Fig. 3 & Supplementary fig. 2). Since inversions may lead to reduced recombination and are known to be instrumental in the emergence of supergenes (Schwander et al., 2014), it is tempting to speculate that a supergene combining several beneficial alleles in relation to insecticide exposure segregate in *C. lectularius* populations. This hypothesis should be further tested in the future.

## 5 LIMITS OF THE PRESENT STUDY AND PERSPECTIVE

In this study, we focused our attention on selected variants that had particularly high frequency in both LF and SF strains compared to LL and GL strains, while taking into account their common history. In species with large (recent) effective population size, the very same mutant may be repeatedly selected in different environments under similar selective pressure, through convergent evolution (Karasov et al., 2010). However, species with lower effective present-day population size are likely limited by the mutation rate, and such convergence is unlikely. It is unclear whether bed bug populations are limited by mutations or not. If they are, then we do not expect similar mutants to be selected in different environments, unless migration from one population brings the favorable allele in the second population. Additionally, since multiple pathways to pyrethroid resistance may evolve in different populations (Djouaka et al., 2008), we acknowledge that this analysis may miss insecticide resistance mutations that would be specific to one or another resistant population.

Another limit of our study relies on the fact that we considered “derived” alleles as opposed to “ancestral”, carried by the Harlan strain from which the reference genome was sequenced. Harlan strain was collected in Fort Dix (New Jersey, USA), back in 1973. We thus tried to identify mutations involved in so-called hard selective sweeps. Many examples of hard sweeps have been described as being involved in insecticide resistance (Cohen et al., 1985; Hawkins et al., 2019; Calla et al., 2021). However, adaptation may also occur through soft sweeps involving standing genetic variation (Hawkins et al., 2019). In that case, the mutants segregated during some time in the population before the selective pressure started. If adaptation to pyrethroids involves standing genetic variation, then our pipeline may have filtered out some valid candidates. However, standing alleles that later increase in frequency due to positive selection are usually at low initial frequency (before the environmental shift). In the three-spined stickleback, where adaptation from standing genetic variation has been extensively studied (DeFaveri and Merilä, 2014; Heckwolf et al., 2020), marine populations adapted repeatedly through standing genetic variation to freshwater habitat. Most adaptive haplotypes in fresh water environment can be found at frequency < 10% in marine environment (Bassham et al., 2018). Since the reference genome has a small probability of having a low frequency allele, then our pipeline still has a high probability of considering it as “derived”. In addition, even in case the Harlan strain carried a later on beneficial allele, our pipeline should at least identify linked “derived” (possibly neutral) variants, initially associated with the beneficial mutant through linkage disequilibrium. In conclusion, we acknowledge that our pipeline may have missed some adaptive alleles from standing genetic variation, although the effect may be limited.

Whether they arose through soft or hard selective sweeps, and whether they involve protein changes or not, it is crucial to identify insecticide resistance mutations in bed bug populations. As our study was restricted to the analysis of intra-scaffold SVs, thus preventing the detection of events spanning the entire superlocus, future efforts should be made to clarify the genomic structure and polymorphism of LG 11. This can be achieved using long read sequencing and/or Hi-C data. This could clarify the structure of the haplotypes and may help to identify the genetic determinants underlying the resistance evolution. From an applied perspective, knowing the genetic basis of such mutations can be used to design diagnostic tests, and thus to track insecticide resistance in the field (Van Leeuwen et al., 2020).

## acknowledgements

This work was performed using the computing facilities of the CC LBBE/PRABI.

## data availability statement

Raw data that support the findings of this study are openly available on SRA under BioProject PRJNA826750. Global SNP pool described in section 2.5.1 is available on Dryad (doi.org/10.5061/dryad.9cnp5hqp6).

