## Supplemental Table 2-4, Fig 1-6 for "A major 6 Mb superlocus is involved in pyrethroid resistance in the common bed bug *Cimex lectularius*"

### 6 | SUPPLEMENTARY

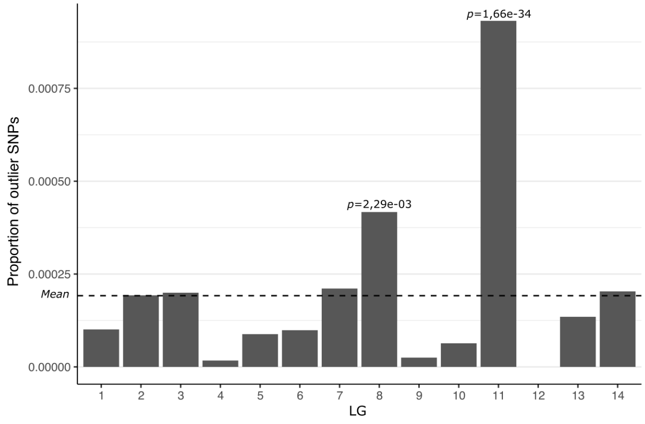


**SUPPLEMENTARY FIGURE 1** Proportions of outlier SNPs among SNPs in linkage groups (LG). The dashed line represents average proportion (a = 0.00019). Binomial test were performed between observed proportion and average proportion in each LG, and significant corrected p-values were detailed above corresponding LG.

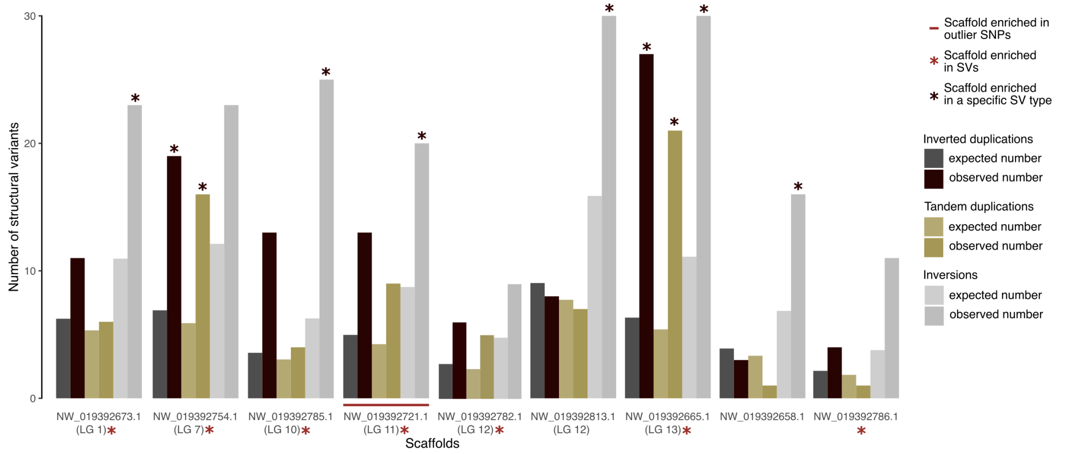


**SUPPLEMENTARY FIGURE 2** Expected and observed numbers of structural variants (SVs) for scaffolds enriched with structural variants in general (delineated with a red star below) or enriched with a specific type of structural variant (namely tandem duplications, inverted duplications and inversions, delineated with a black star above corresponding observed number). The scaffold also enriched with outlier SNPs is underlined in red.

| | $n$ | slope | $df$ | $X^2$ | $LD_{95}$ | 95% mortality CI95 |
|---|---|---|---|---|---|---|
| London Lab | 280 | 1.840 | 27 | 178.10 | 1.3 | 91.2-98.8% |
| German Lab | 200 | 3.123 | 19 | 172.97 | 0.4 | 89.8-100% |
| London Field | 170 | 1.030 | 17 | 75.76 | 21.0 | 89.7-100% |
| Sweden Field | 210 | 0.941 | 21 | 97.04 | 69.2 | 90.5-99.5% |

**SUPPLEMENTARY TABLE 2** Additional information for dose-mortality analysis performed on *Cimex lectularius* strains. Number of insects ($n$), slope, degree of freedom ($df$) and chi-square ($X^2$, computed as null deviance minus residual deviance), lethal dose (in ng) for 95% of insects in the population ($LD_{95}$) were given for each strains. The confidence interval at 95% (CI95) for 95% mortality was also computed for each strain using the predict function from the R/ecotox package.

| Scaff. | Gene ID | Position | Annotation | Location | $F_{alt}$ LL | $F_{alt}$ LF | $F_{ST}$ (pval) |
|---|---|---|---|---|---|---|---|
| 2763 | 106666926 | 419008 | *GST1-1*-like | exon NS | 0.20 | 0.38 | -0.01 (0.38) |
| | 106666926 | 419242 | *GST1-1*-like | exon NS | 0.08 | 0.27 | 0.05 (0.25) |
| | 106666926 | 419369 | *GST1-1*-like | exon NS | 0.06 | 0.24 | 0.03 (0.28) |
| | 106663963 | 503362 | *GST*-like | exon NS | 0.10 | 0.46 | 0.16 (0.10) |
| 2721 | 106663983 | 1833078 | *CYP6B5*-like | exon NS | 0.50 | 0.93 | 0.29 (0.03*) |
| | 106663983 | 1833420 | *CYP6B5*-like | exon NS | 0.41 | 0.79 | 0.17 (0.09) |
| | 106663983 | 1833792 | *CYP6B5*-like | exon NS | 0.46 | 0.78 | 0.11 (0.15) |
| | 106663983 | 1833793 | *CYP6B5*-like | exon NS | 0.46 | 0.76 | 0.09 (0.18) |
| | 106663983 | 1833810 | *CYP6B5*-like | exon NS | 0.50 | 0.77 | 0.07 (0.20) |
| | 106663983 | 1833837 | *CYP6B5*-like | exon NS | 0.56 | 0.75 | -0.02 (0.40) |
| | 106663983 | 1836843 | *CYP6B5*-like | exon NS | 0.58 | 1,00 | 0.33 (0.02*) |
| | 106663983 | 1838158 | *CYP6B5*-like | exon NS | 0.55 | 0.85 | 0.13 (0.12) |
| | 106663982 | 1849007 | *CYP6B1*-like | exon NS | 0.27 | 0.72 | 0.28 (0.04*) |
| | 106663982 | 1849044 | *CYP6B1*-like | exon NS | 0.29 | 0.75 | 0.2 (0.03*) |
| | 106663982 | 1849067 | *CYP6B1*-like | exon NS | 0.29 | 0.63 | 0.15 (0.11) |
| | 106663982 | 1849919 | *CYP6B1*-like | exon NS | 0.59 | 0.84 | 0.07 (0.21) |
| | 106663981 | 1873544 | *CYP6B1*-like | exon NS | 0.33 | 0.50 | 0.00 (0.35) |
| | 106663981 | 1873558 | *CYP6B1*-like | exon NS | 0.33 | 0.53 | 0.02 (0.30) |
| | 106663981 | 1874026 | *CYP6B1*-like | exon NS | 0.59 | 0.66 | -0.05 (0.50) |
| | 106663981 | 1874033 | *CYP6B1*-like | exon NS | 0.54 | 0.57 | -0.06 (0.53) |
| | 106663981 | 1875827 | *CYP6B1*-like | exon NS | 0.07 | 0.25 | 0.01 (0.32) |
| | 106663981 | 1875904 | *CYP6B1*-like | exon NS | 0.23 | 0.35 | -0.04 (0.46) |
| | 106663981 | 1875976 | *CYP6B1*-like | exon NS | 0.72 | 0.75 | -0.09 (0.69) |

**SUPPLEMENTARY TABLE 3** Non-synonymous (NS) SNPs falling within putative resistant genes inside the superlocus detected, with frequency for derived allele from ancestral susceptible population ($F_{alt}$) superior in London Field (London Field) than in London Lab (London Lab). Gene annotation *GST* stands for glutathione-S-transferase, *CYP* for cytochromes P450. To obtain scaffold name, one should add "NW_01939" at the beginning, and ".1" at the end.

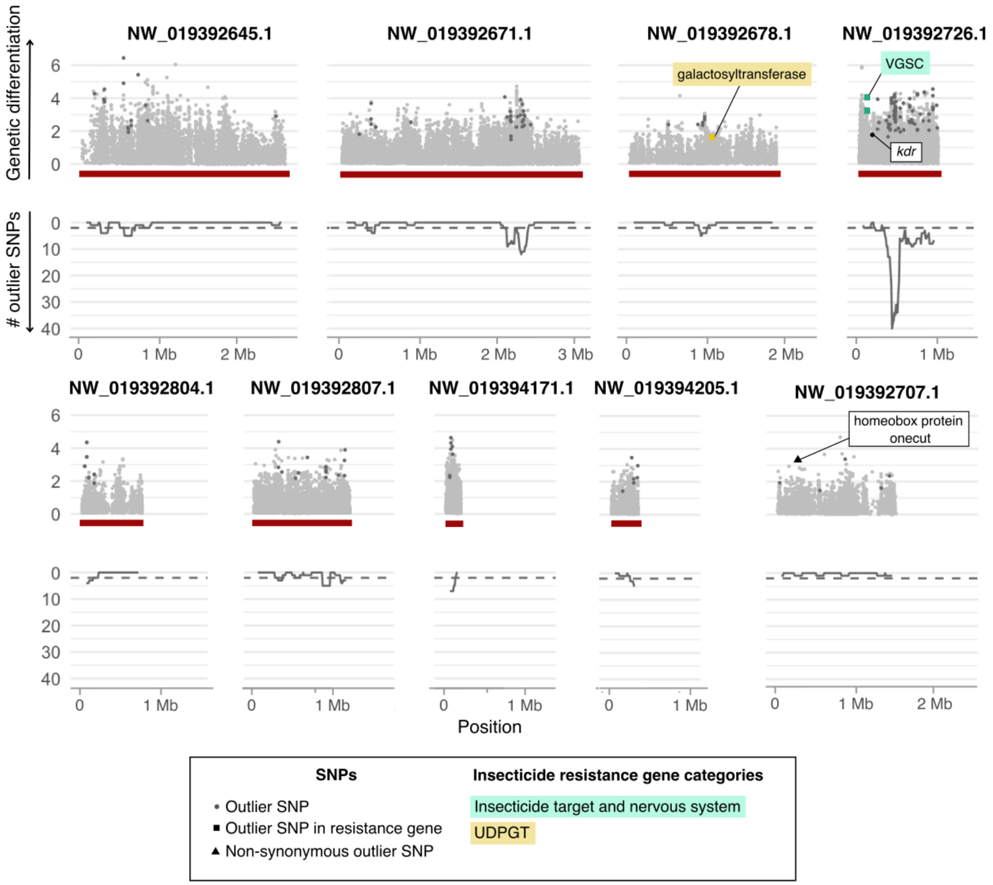


**SUPPLEMENTARY FIGURE 3** Several supplementary scaffolds for *Cimex lectularius* genome not located on linkage groups. Scaffolds detected as significantly enriched in outlier SNPs are underlined in red. The scaffold carrying the unique outlier non-synonymous SNP is also represented (NW_019392707.1). In the upper part of a plot, the empirical p-value ($p$) of genetic differentiation index ($F_{ST}$) between London Lab and London Field populations in -log10 scale is plotted. The *x*-axis indicates the position within each scaffold. Outlier SNPs (see Materials & methods) are distinguished by a darker grey. Square-shaped points are the outlier SNPs that fall within candidate resistance genes, and are colored according to those genes' categories. Their protein products are labeled. Non-synonymous outlier mutation was represented with a black triangle. Although not an outlier SNP, the location of the non-synonymous *kdr* L925I mutation was given as an indication. Positions of genes with an expected amplified copy number are displayed with a dashed black vertical line and labelled likewise. In the lower part, "# outlier SNPs" represents the number of outlier SNPs counted in each 100 kb sliding window with a step of 10 kb. Dashed line stands for a threshold of 2 outlier SNPs, which delineates quantile 99%.

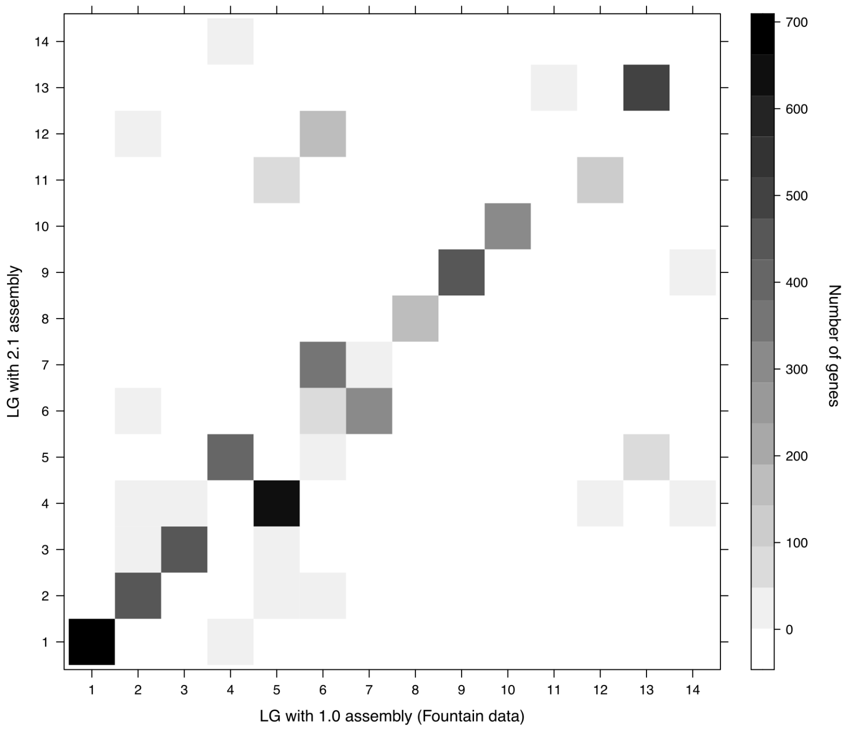


**SUPPLEMENTARY FIGURE 4** Joint-distribution of genes in Linkage Groups (LG) in Fountain et al. (2016) paper and the present paper. Genes present on the first diagonal are located on the same LG in both cases.

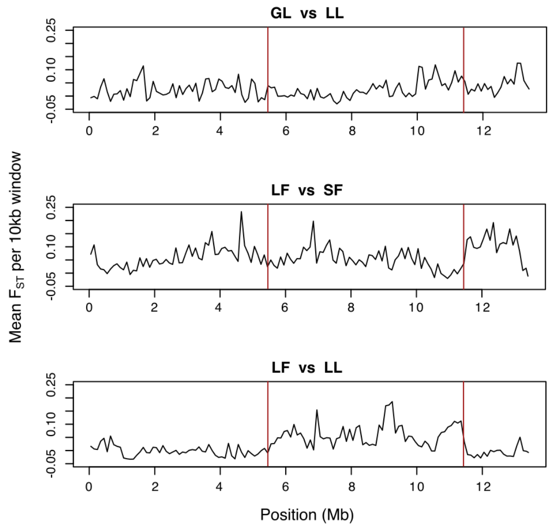


**SUPPLEMENTARY FIGURE 5** Differentiation between strains over LG 11. Mean $F_{ST}$ values were computed in 10 kb windows with 10 kb step for three comparisons: between the two susceptible strains (GL = German Lab and LL = London Lab), the two resistant strains (LF = London Field and SF = Sweden Field), and between Londonian strains (LF and LL). The position of a "superlocus" (NW_019392721.1, NW_019392763.1 and NW_019942502.1) is delineated with red lines.

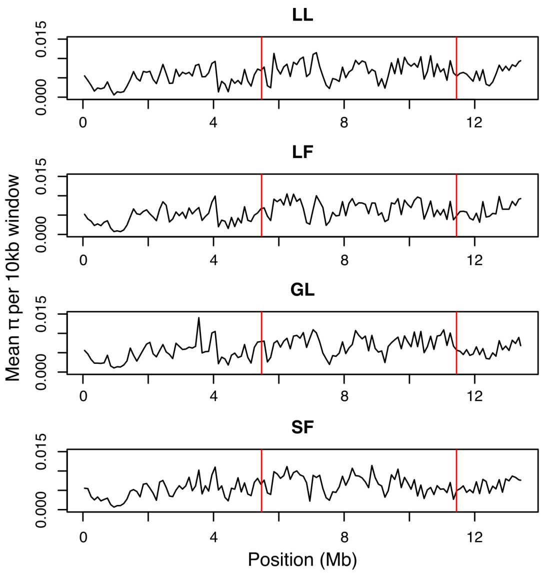


**SUPPLEMENTARY FIGURE 6** Nucleotidic diversity within strains over LG 11. Mean $\pi$ values were computed in 10 kb windows with 10 kb step for the four strains: two susceptible (GL = German Lab and LL = London Lab) and two resistant (LF = London Field and SF = Sweden Field). The position of a "superlocus" (NW_019392721.1, NW_019392763.1 and NW_019942502.1) is delineated with red lines.

| LG | Scaff. | Event | Size | Gene ID | Annotation | *F* LL | *F* LF | $F_{ST}$ | RDR (q) |
|---|---|---|---|---|---|---|---|---|---|
| 1 | 2640 | Inv | 324 kb | 106672175 | sulfotransferase | 0 | 0.09 | 0.05 (7%) | 1.23 (1.21-1.24) |
| 1 | 2679 | InvDup | 1.494 Mb | 106661337 | sulfotransferase | 0.06 | 0.24 | 0.07 (4%) | 1.23 (1.23) |
| | | | | 106661366 | choline kinase | | | | |
| | | | | 106661634 | CYP450 306a1 | | | | |
| | | | | 106661588 | CYP450 18a1 | | | | |
| 1 | 2641 | Inv | 262 kb | 106669934 | SOD [Cu-Zn] | 0 | 0.10 | 0.05 (6%) | 1.22 (1.21-1.24) |
| | | | | 106669936 | SOD [Cu-Zn] 2 | | | | |
| 2 | 2979 | Inv | 7 kb | 106665265 | endocuticle str. prot. | 0 | 0.12 | 0.06 (4%) | 1.15 (1.15-1.29) |
| | | | | 106665194 | endocuticle str. prot. | | | | |
| 13 | 2685 | Inv | 76 kb | 106662993 | cuticle protein | 0.16 | 0.41 | 0.08 (2%) | 1.23 (1.20-1.25) |
| | | Inv | 72 kb | 106662993 | cuticle protein | 0 | 0.09 | 0.05 (7%) | 1.24 (1.20-1.25) |
| - | 2647 | Inv | 484 kb | 106668191 | NADPH oxidored. | 0.06 | 0.26 | 0.08 (3%) | 1.23 (1.23) |
| - | 2656 | InvDup | 381 kb | 106669691 | glucosyltransf. | 0.22 | 0.47 | 0.06 (4%) | 1.24 (1.23) |
| - | | Inv | 221 kb | 112126170 | galactosyltransf. | 0 | 0.24 | 0.13 (1%) | 1.22 (1.21-1.24) |
| | | | | 106669680 | glucuronosyltransf. | | | | |
| - | 2881 | InvDup | 27 kb | 106665141 | cuticle protein | 0.02 | 0.16 | 0.06 (5%) | 1.29 (1.26) |
| | | | | 106664994 | cuticle protein | | | | |
| | | | | 106664995 | cuticle protein | | | | |
| | | | | 106664992 | cuticle protein | | | | |

**SUPPLEMENTARY TABLE 4** Resistance genes overlapping with structural variants with extreme $F_{ST}$ value (top 10%, specified in brackets). SVs events are delineated as tandem duplications (TandDup), inverted duplications (InvDup), and inversions (Inv). Read Depth Ratio (RDR) between London Field and London Lab was given for each event, along with the threshold quantile (q). Since duplications were all detected in higher frequency in London Field, we only showed the q75 quantile used. For inversions, we showed the q25-q75 interval. "SOD" stands for superoxide dismutase. Gene annotations have been shortened, but full informations can be found using gene ID. To obtain scaffold name, one should add "NW_01939" at the beginning, and ".1" at the end.
